# Modelling the distribution of white matter hyperintensities due to ageing on MRI images using Bayesian inference

**DOI:** 10.1101/327205

**Authors:** Vaanathi Sundaresan, Ludovica Griffanti, Petya Kindalova, Fidel Alfaro-Almagro, Giovanna Zamboni, Peter M. Rothwell, Thomas E. Nichols, Mark Jenkinson

## Abstract

White matter hyperintensities (WMH), also known as white matter lesions, are localised white matter areas that appear hyperintense on MRI scans. WMH commonly occur in the ageing population, and are often associated with several factors such as cognitive disorders, cardiovascular risk factors, cerebrovascular and neurodegenerative diseases. Despite the fact that some links between lesion location and parametric factors such as age have already been established, the relationship between voxel-wise spatial distribution of lesions and these factors is not yet well understood. Hence, it would be of clinical importance to model the distribution of lesions at the population-level and quantitatively analyse the effect of various factors on the lesion distribution model.

In this work we compare various methods, including our proposed method, to generate voxel-wise distributions of WMH within a population with respect to various factors. Our proposed Bayesian spline method models the spatio-temporal distribution of WMH with respect to a parametric factor of interest, in this case age, within a population. Our probabilistic model takes as input the lesion segmentation binary maps of subjects belonging to various age groups and provides a population-level parametric lesion probability map as output. We used a spline representation to ensure a degree of smoothness in space and the dimension associated with the parameter, and formulated our model using a Bayesian framework.

We tested our algorithm output on simulated data and compared our results with those obtained using various existing methods with different levels of algorithmic and computational complexity. We then compared the better performing methods on a real dataset, consisting of 1000 subjects of the UK Biobank, divided in two groups based on hypertension diagnosis. Finally, we applied our method on a clinical dataset of patients with vascular disease.

On simulated dataset, the results from our algorithm showed a mean square error (MSE) value of 7.27 × 10^−5^, which was lower than the MSE value reported in the literature, with the advantage of being robust and computationally efficient. In the UK Biobank data, we found that the lesion probabilities are higher for the hypertension group compared to the non-hypertension group and further verified this finding using a statistical t-test. Finally, when applying our method on patients with vascular disease, we observed that the overall probability of lesions is significantly higher in later age groups, which is in line with the current literature.

## 1. Introduction

White matter hyperintensities of presumed vascular origin (WMH), also called white matter lesions (Wardlaw et al. (2013)), are common findings in the brain white matter that appear hyperintense on T2-weighted, fluid attenuated inversion recovery (FLAIR), and proton density-weighted brain MRI images. Even though the pathogenesis of WMH has not yet been well understood (Wardlaw et al. (2013)), WMH are strongly associated with cerebrovascular disease and vascular risk factors (Li et al. (2013)), and they are also frequently found in neurodegenerative diseases such as Alzheimers disease (Debette and Markus (2010), Prins and Scheltens (2015)). In the general population, they have been associated with increased risk of stroke, dementia and death (Debette et al. (2010)).

Several visual rating scales for WMH are available and commonly used (Wahlund et al. (2001), Fazekas et al. (1987)). However, determining the voxel-wise distribution of WMH at the population-level is important in order to investigate the relationship between the spatial distribution of lesions and various factors such as age, vascular risk factors and cognitive function. This would aid in studying differences in lesion distribution between normal and pathological ageing (Biesbroek et al. (2013), Duering et al. (2014)). In fact, although the occurrence of WMH is common in the ageing population, clear relationships between amount and distribution of lesions and the cognitive, demographical, and risk factors is not yet well understood.

The simplest method used in the literature for correlating voxel-wise lesion distribution maps with various parametric factors (such as age, cognitive function and so on) includes averaging subject-level lesion probability maps (Rostrup et al. (2012)) that are grouped according to those parametric factors. However, this approach does not necessarily provide continuity or smoothness in space or across the parametric dimension for the lesion probabilities. Alternatively, several *mass-univariate* methods such as voxel-based lesion-symptom mapping (VLSM) (Bates et al. (2003)) and voxel-wise linear regression of lesion probabilities against various factors (Charil et al. (2003), Charil et al. (2007)) have been proposed. For example, Charil et al. (2007) studied the correlation between the cortical thickness and various factors including lesion distribution in multiple sclerosis using voxel-wise linear regression. However, these methods are based on models that work independently in each voxel and do not capture the relationship between neighbouring voxels. Hence, they are not ideal for modelling the lesion distribution, since lesions are clustered regions rather than isolated voxels. Moreover, the linear models used in these methods are not optimal for binary data (Charil et al. (2007), Bates et al. (2003)).

Spatially varying coefficient processes establish a local spatial relationship between the coefficients of regression models (Gelfand et al. (2003); Gamerman et al. (2003); Ge et al. (2014)). For example, Bayesian Spatial Generalized Linear Mixed Model (BSGLMM) proposed by Ge et al. (2014) is based on spatially varying coefficients to determine the relationship between the spatial distribution of lesions and subject specific covariates such as multiple sclerosis (MS) subtype, age, gender, disease duration and disease severity measures. The spatially varying coefficients are modelled jointly using a multivariate pairwise difference prior model, a particular instance of the Multivariate Conditional Autoregressive model. However, the dimension of the correlation matrices involved in spatially varying coefficient processes is very high and their inversion becomes computationally infeasible for very large imaging datasets (Ge et al. (2014)). In fact, the computational load of the model proposed by Ge et al. (2014) requires the use of graphical processing units for parallel computing.

Several solutions have been proposed to overcome this problem, including Gaussian predictive processes (Banerjee et al. (2008)), using a truncated Karhunen-Love expansion to estimate spatially varying coefficients (Crainiceanu et al. (2009)), functional principal component analysis approach (Reiss and Ogden (2010)) and covariance tapering (Kaufman et al. (2008)). However, all of them perform data reduction (unlike Ge et al. (2014)) and depend on strict assumptions (Crainiceanu et al. (2009)).

In this paper, we propose an alternative approach by developing a probabilistic model to obtain a parametric lesion probability map in order to investigate the relationship between distribution of WMH and various parametric factors. Our method is not computationally intensive, and uses a spline representation to ensure continuity and smoothness between neighbouring voxels in both spatial and parametric dimensions. In our model, the lesion distribution at each voxel is obtained by the linear combination of all the splines on the neighbouring voxels. Moreover, our model adopts a Bayesian framework in order to overcome the discretely sampled nature of the input binary maps obtained from lesion segmentation. Our approach for modelling the spatial distribution of lesions is suitable for very large imaging datasets such as UK Biobank (Miller et al. (2016)) and allows the flexibility of observing the effect of various parametric factors on the lesion distribution. Although in principle our model could work with any factor, in this work we analysed the voxel-wise distribution of lesions with respect to age since a strong relationship between the progression of WMH and age has already been established (Simoni et al. (2012)).

We first evaluated our model on a simulated dataset and compared it to existing methods. We then validated our model on two real datasets. The first one is a sub sample of the UK Biobank, in which we tested the relationship between WMH and age comparing subjects with and without hypertension. We chose hypertension as our grouping variable, since it has been found to be one of the important risk factors of WMH (Dufouil et al. (2001), Gottesman et al. (2010)) in addition to age. Also in this case we compared our results against the existing methods. As a final validation, we applied our model to a clinical population (vascular population of subjects who had a transient ischemic attack or minor stroke) and analysed the results with respect to age.

## 2. Methods

### 2.1. Modelling the distribution of WMH using Bayesian inference

Our proposed Bayesian spline model takes as input the binary lesion map and the age (or other parametric factor of interest) for each individual subject. We model the distribution of lesions in three steps: 1) constructing a representation for the lesion probability distribution using by spline basis functions, 2) formulation of the posterior lesion probability; and 3) maximisation of the posterior probability by constrained optimisation.

Let *D*_*s*_ be the 3D binary lesion map of individual subject *S*. Our aim was to form a 4D spatio-temporal parametric lesion probability map with the 4^*th*^ dimension indicating the parametric factor of interest.

#### 2.1.1. Constructing a representation for the lesion probability distribution using by spline basis functions

Since the binary lesion maps contains localised areas, the lesion voxels are sparse and have discrete values (0 or 1). Therefore, to ensure spatial and temporal continuity in the probability distribution of lesions, we approximate the lesion probabilities using cubic b-splines.

Consider cubic b-splines with spline coefficients *C*_*i*_ and basis functions *B*_*ij*_, where *i* denotes the indices of knot points of the basis functions and *j* = (*x,y,z,t*) indicate the spatial coordinates in the first 3 dimensions and the parametric factor in the 4^*th*^ dimension. The lesion probability *θ*_*j*_, at each voxel is related to *C*_*i*_ and *B*_*ij*_, by Eqn. 1:

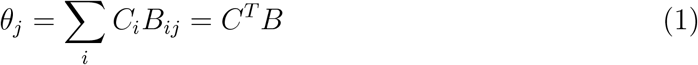

The probability value at each voxel is calculated as the linear combination of spline basis functions. We formulated the splines as a separable outer product of individual 1D splines in 4 dimensions (3 in space and one in the parametric dimension). To implement the above calculation, we generated 1-dimensional cubic b-splines and convolved the binary map with scaled 1D b-splines independently in all 4 dimensions to get spatially continuous lesion probabilities *θ* as shown in Figure 1. This operation is more computationally efficient than off-the-shelf spline fitting toolboxes and is well suited to large datasets.

It is worth noting that after each convolution edge artefacts occurred due to convolution with the tails of the splines and zero padding. In order to get the corrected convolution output *f*_*norm*_, we divided the convolution output *f*_*in*_ with a normalizing image (formed by convolving the same b-splines *B* with an unity image *u*(*x*), where *u*(*x*) = 1 inside the valid FOV - brain voxels - and 0 outside, thus forming a similar pattern at the edges) as shown in eqn. 2.

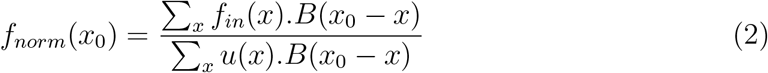

**Figure 1:**
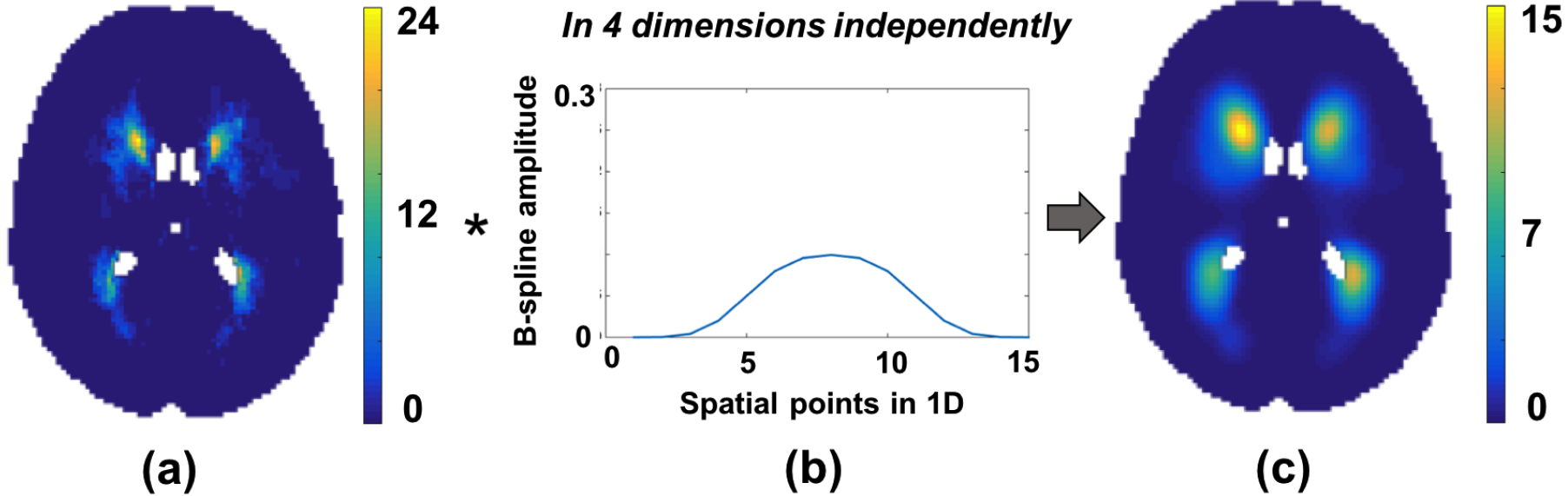
Spline modelling with cubic b-splines. (a) Average of binary lesion maps that are grouped according to a parametric factor, (b) Cubic b-spline used for modelling, (c) Spatially continuous lesion probability map obtained as a result of spline fitting. The convolution of (b) with (a) is done in all 4 dimensions independently to get (c).

#### 2.1.2. Formulation of the posterior lesion probability

The most common lesion probability estimation method is averaging lesion maps across subjects. However, the accuracy of this approach is sensitive to the amount of data, especially considering the fact that there might not be any subject representing specific age groups or a few subjects might not have any WMH. Bayesian methods are well suited to this problem since they allow for the uncertainty associated with the limited amount of data, while the spline model ensures continuity between the neighbouring voxels.

As we consider age as our parametric factor, the 4^*th*^ dimension, *t*, of our data indicates the bin number corresponding to the age groups. We formed these bins *t* by grouping the subjects into age groups *age*_*t*_ (e.g., *age*_1_ = (20, 22], *age*_2_ = (23, 25], etc.). The age groups can be defined to have a duration as short as we require (even in months) and hence binning the ages does not necessarily restrict the values to be overly discretised in the parametric dimension. Let *R* (shown in Figure. 2(a)) be the 4D spatio-temporal volume formed by summing the binary lesion maps of the subjects in each age group *age*_*t*_. The value at each voxel *R*_*j*_ = *R*(*x,y,z,t*) denotes the number of subjects having lesions at that voxel in the age group *age*_*t*_. Let *N* be a 4D volume in which the value at each voxel *N*_*j*_ = *N*(*x,y,z,t*) denotes the number of subjects in each age group *age*_*t*_.

**Figure 2:**
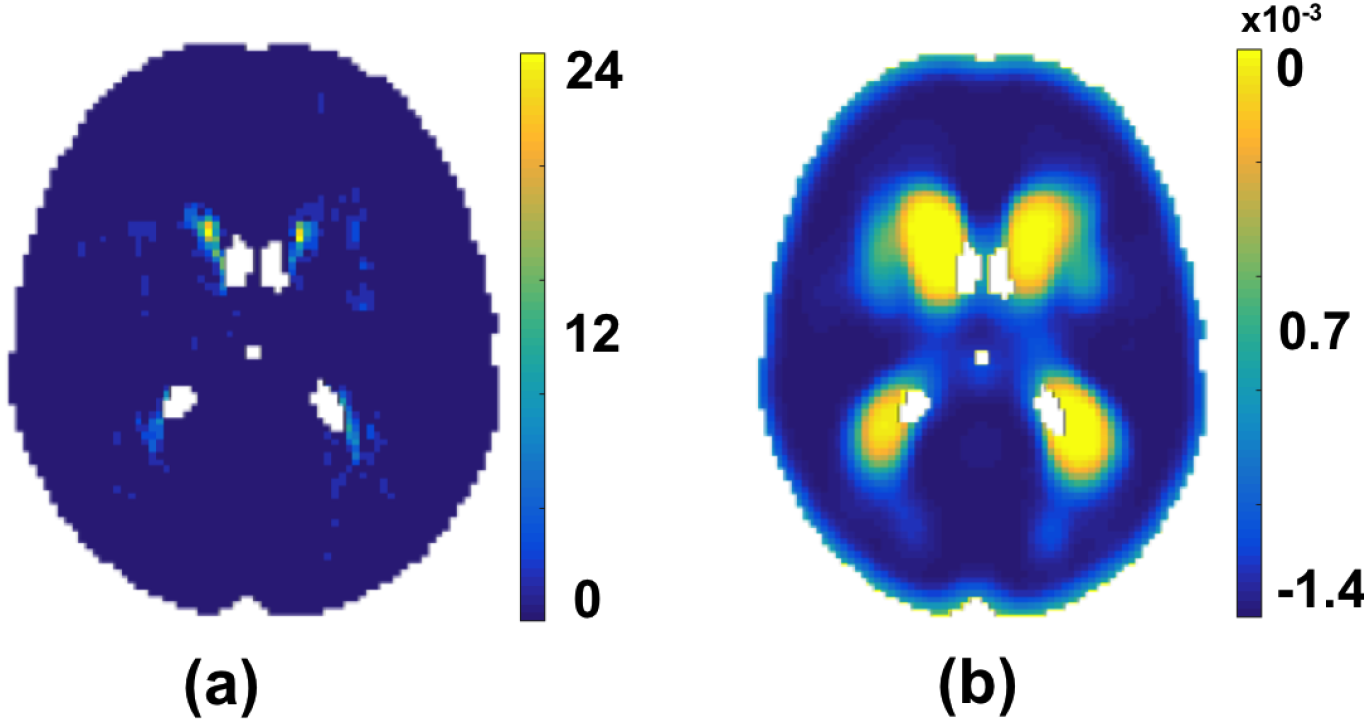
Accumulation of data. (a) In each age group *age*_*t*_, the number of subjects having lesions (*R*_*j*_). (b) Gradient of the log-probability distribution obtained by Eqn. 16

*R* and *N* are formed by,

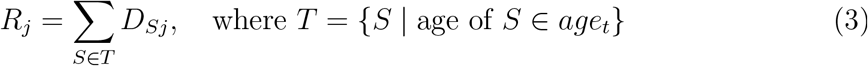

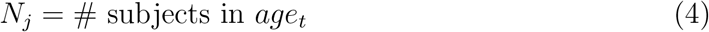

For an age group *age*_*t*_, *R*_*j*_ indicates number of observations of binary data (lesion occurrence) in the subjects belonging to this age group *age*_*t*_, with the total number of subjects in *age*_*t*_ given by *N*_*j*_. Thus the probability of observing *R*_*j*_ number of binary outputs given *N*_*j*_ and probability of lesion occurrence *θ*_*j*_ for an age group *age*_*t*_ is given by the binomial likelihood distribution,

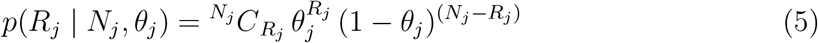

where 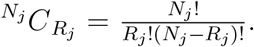 The full likelihood distribution (assuming independence between age groups and voxel locations) for all age groups and at all voxel locations is given by the product of individual likelihoods,

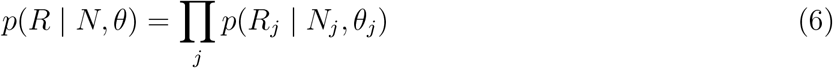

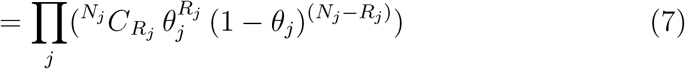

Using Bayes theorem, the posterior lesion probability distribution *p*(*θ* | *N*, *R*) is given by,

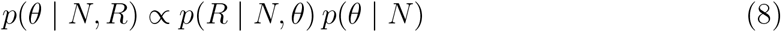

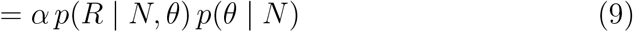

We will assume that the prior probability of lesion occurrence is the same for any point in space and time. Hence with uniform prior *p*(*θ*) = 1,

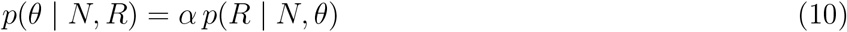

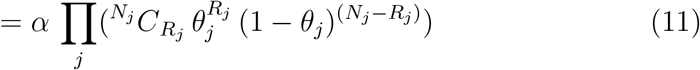

Our aim is to maximize *p*(*θ* | *N*, *R*) to obtain a parametric model of the lesion probability distribution over a population. To make the calculation of derivatives simpler, we determine the log-posterior function since the logarithm is a monotonically increasing function. Log-posterior *L*(*θ* | *N*, *R*) is given by,

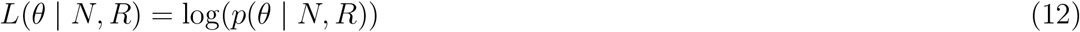

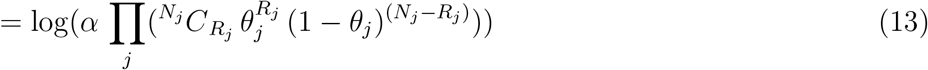

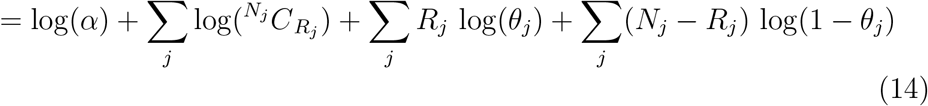

Since the lesion probability values *θ* have been approximated with spline basis functions as explained in the section above, we substitute the values of *θ* from the eqn. 1 in the above log-posterior equation,

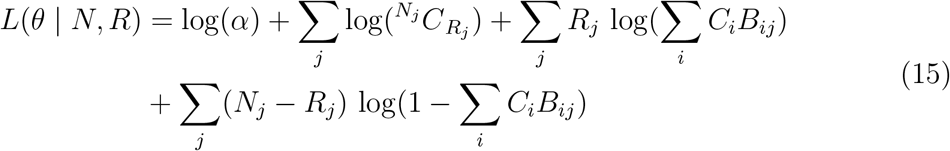

The derivative of *L*(*θ*|*N*, *R*) (shown in Figure. 2(c)) with respect to the spline coefficients *C*_*k*_ is given by,

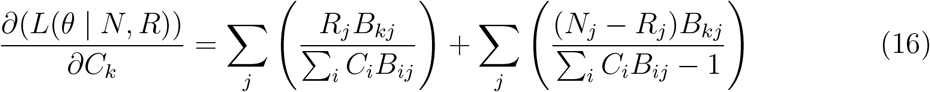

Note that the derivative function requires more spline modelling steps (basically, each summation requires a 4D spline approximation), however, performing 1D convolutions independently in 4 dimensions considerably reduces the running time.

#### 2.1.3. Maximisation of the posterior probability by constrained optimisation

In order to estimate the final lesion probability *θ*_*i*_, we need to determine the spline coefficient values that would maximise the log-posterior. Also, by maintaining the value of *C*_*i*_ within the range [0,1] the final *θ*_*i*_ values will be constrained to the range [0,1]. Hence, to determine the value of *C*_*i*_ corresponding to the *maximum a posteriori* estimate, we formulated it as a constrained optimisation problem as specified below:

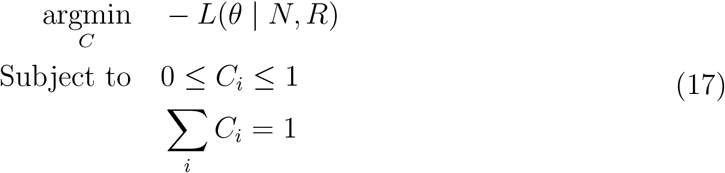

We used the steepest gradient descent algorithm, which is a first order optimisation algorithm that uses an iterative polynomial line search method for the above optimisation. The step size γ for each iteration is determined by a polynomial line search method satisfying Armijo - Wolfe conditions (Wolfe (1969)). In our case, the function to be minimised is –*L*(*θ* | *N*, *R*). For *L*: ℝ^*n*^ → ℝ, the step size for each iteration follows the following condition:

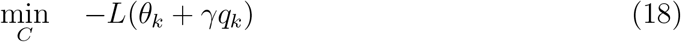

where, *θ*_*k*_ is the current guess, *q*_*k*_ indicates the search direction and γ is the step size. Moreover, in order to speed up the convergence, we redefined *C*_*i*_ using a normalising function (with parameter λ_*i*_ ∈ (–∞, +∞)) in order to maintain the range of *C*_*i*_ within [0,1] and relax the optimisation constraints as given by,

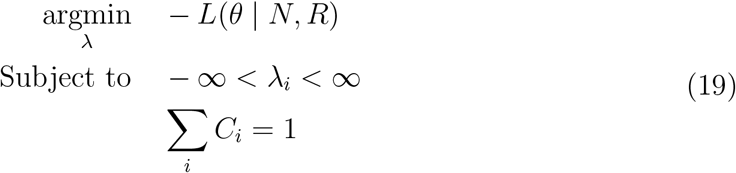

where *C*_*i*_ = (1 + tanh(λ_*i*_))/2 so that *C*_*i*_ ∈ [0,1] when λ_*i*_ ∈ (–∞, +∞). Using the chain rule, the derivative of *L*(*θ*|*N*,*R*) (eqn(16)) can be modified as

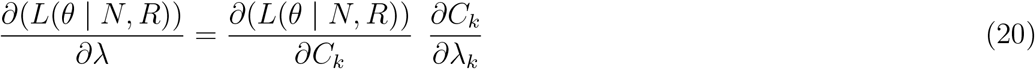

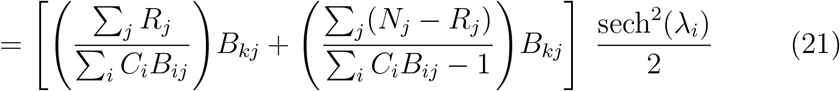

We perform a step of spline modelling (given by eqn. 1) on the optimal *C*_*i*_ that maximises the log-posterior –*L*(*θ* | *N*, *R*) to get the final estimated parametric lesion probability map 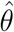. We determined the average of lesion binary maps of subjects within the age group *age*_*t*_ to get the age group-wise average 4D lesion map *Average* = ∑_*j*_(*R*_*j*_/*N*_*j*_). We provided smoothed *Average* as the initial estimate for *C*_*i*_ during optimisation.

Figure 3 illustrates the initial and final values of *C*_*i*_ and λ_*i*_ involved in the optimization step for two different age groups (taken from different age bins of the 4D spatio-temporal map). The initial *θ* values (fig. 3(d)) and the final 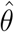 values (fig. 3(e)) are shown with their difference (fig. 3(f)). The results show little difference between the initial *θ* values and final 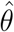 values for the younger age group (≤ 0.005 in fig. 3(f, bottom row)) and a higher difference (≈ 0.01 in fig. 3(f, top row)) in the elder age group. This could be attributed to the difference in the amount of lesion voxels between the age groups.

**Figure 3:**
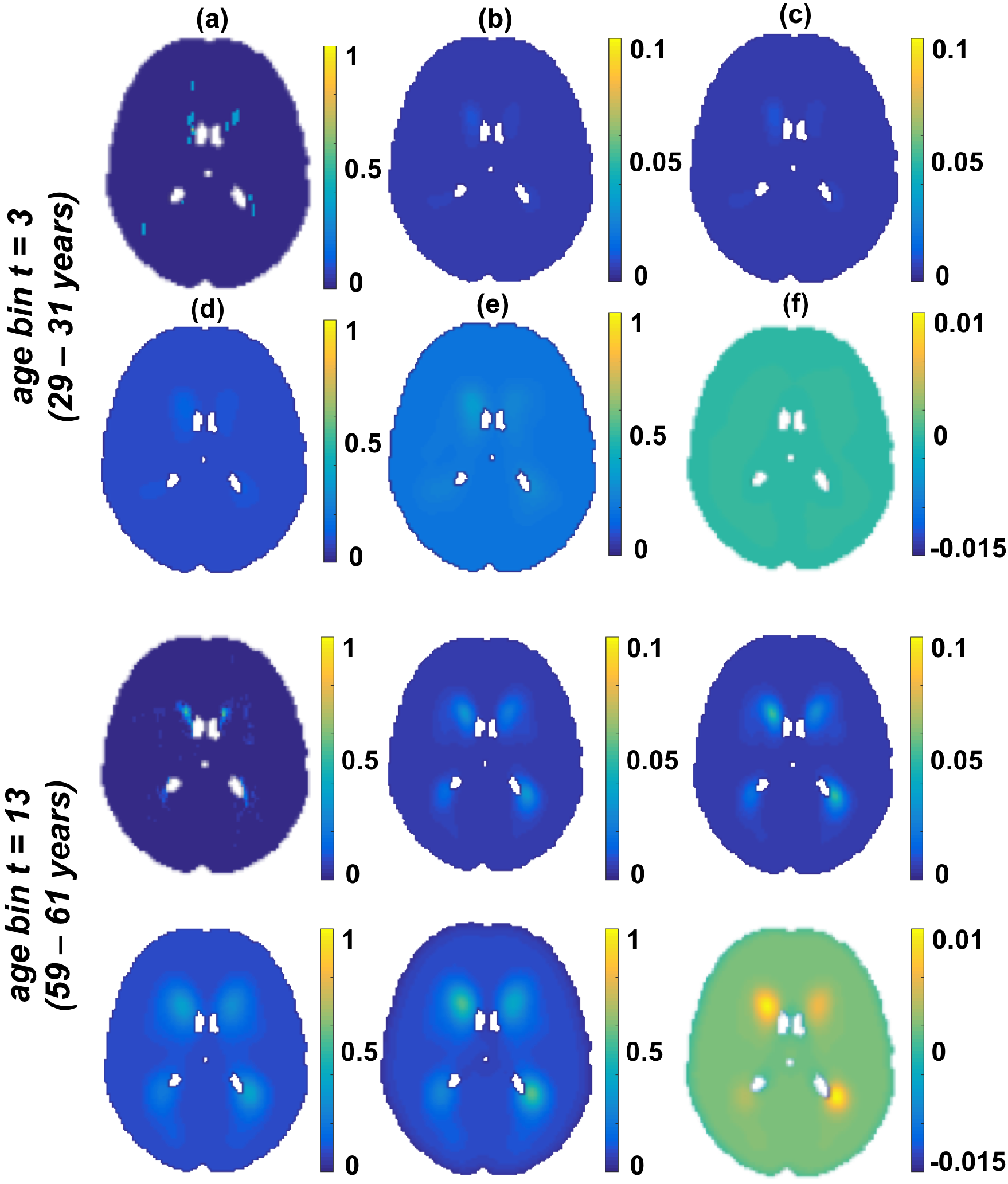
An instance of initial and final values in the optimisation step. The instances relate to two different age groups (younger age group in top rows). All the images are shown at slice *z* = 45. (a) Average of lesions across the age group (*R*_*j*_/*N*_*j*_). (b) Initial *C* values obtained by smoothing *R*_*j*_/*N*_*j*_, (c) Final *C* values after convergence, (d) *θ* values from initial *C* values, (e) 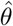 values estimated from final *C* values after convergence. (f) Difference between initial *θ* values and final 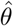 values. Note that in the younger age group there is not much difference (top row) in *θ* before (d) and after (e) convergence, since the neighbouring voxels are mostly non-lesions in the young population, whereas 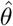 values are larger and differ more in the bottom row between (d) and (e) as shown in (f) in older population.

#### 2.1.4. Convergence analysis

Implementation of steepest gradient descent optimisation algorithm was done in MATLAB (Bortz and Kelley (1998)). We specified the initial step-size γ to be 1 × 10^−3^, while step sizes for the subsequent iterations were determined by the polynomial line search algorithm, for which lower and upper bounds were set to 0.1 × initial γ and 0.5 × initial γ respectively. We specified the maximum number of iterations as 50. The main parameter for determining γ for each iteration is the 2-norm ‖𝒢‖ of the 4D gradient matrix 𝒢 (calculated from eqn. 20). We set the final convergence tolerance value for the decrease in ‖𝒢‖ as 1 × 10^−4^ (since gradient of the function must be zero at the function minimum).

### 2.2. Simulations

In order to evaluate the accuracy of our model, we initially tested it on a simulated dataset. To this aim, we compared the ground truth distribution (*θ*_*true*_) with the probability distribution 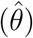 estimated by our algorithm and compared it with other methods.

We simulated volumes of size 91 × 109 × 91 arranged in 6 months age bins to form 60 age groups, thus creating a simulated set of subject images with dimensions 91 × 109 × 91 × 60. For this simulation, we assigned a random number of subjects in each age group *N*_*sim*_ ∈ [1, 50]. For each subject, we generated a lesion map by randomly sampling voxels within a specified ROI (in our case, the MNI brain mask). For sampling, we used the smoothed average of lesion probability maps from a previous study (vascular cohort in Griffanti et al. (2016)) as the ground truth probabilities to obtain a realistic WMH spatial distribution. The probability of sampling each lesion voxel in the data is initially represented by a random normal variable *N*(*μ*(*x*), 1). In order to avoid isolated lesion voxels and to establish spatial dependencies between the neighbouring voxels (so that sampled voxels would look like plausible lesions), we smoothed the sampled lesion voxels using a Gaussian kernel *K* with a standard deviation of 0.8. We obtained binary lesion maps by thresholding the smoothed lesion map *K* ∗ *N*(*μ*(*x*), 1) above 0. As a result of convolution with *K*, the ground truth distribution at voxel *x* is now represented by *N*(*K* ∗ *μ*, *K*^2^ ∗ *M*), where *M* is a binary brain mask. We obtained the true lesion probability for the simulation, *θ*_*true*_, by calculating the cumulative distribution function of the normal variate 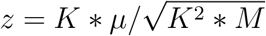 or more specifically, *p*(*z* > 0).

Repeating the above process *N*_*sim*_ times for each age group, we accumulated the sum of binary volumes as *R*_*sim*_ and applied our algorithm to *R*_*sim*_ and *N*_*sim*_ to model the distribution of lesions over the simulated population, to get 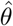 (algorithm output). An instance of data obtained from the simulation is shown in Figure 4.

**Figure 4:**
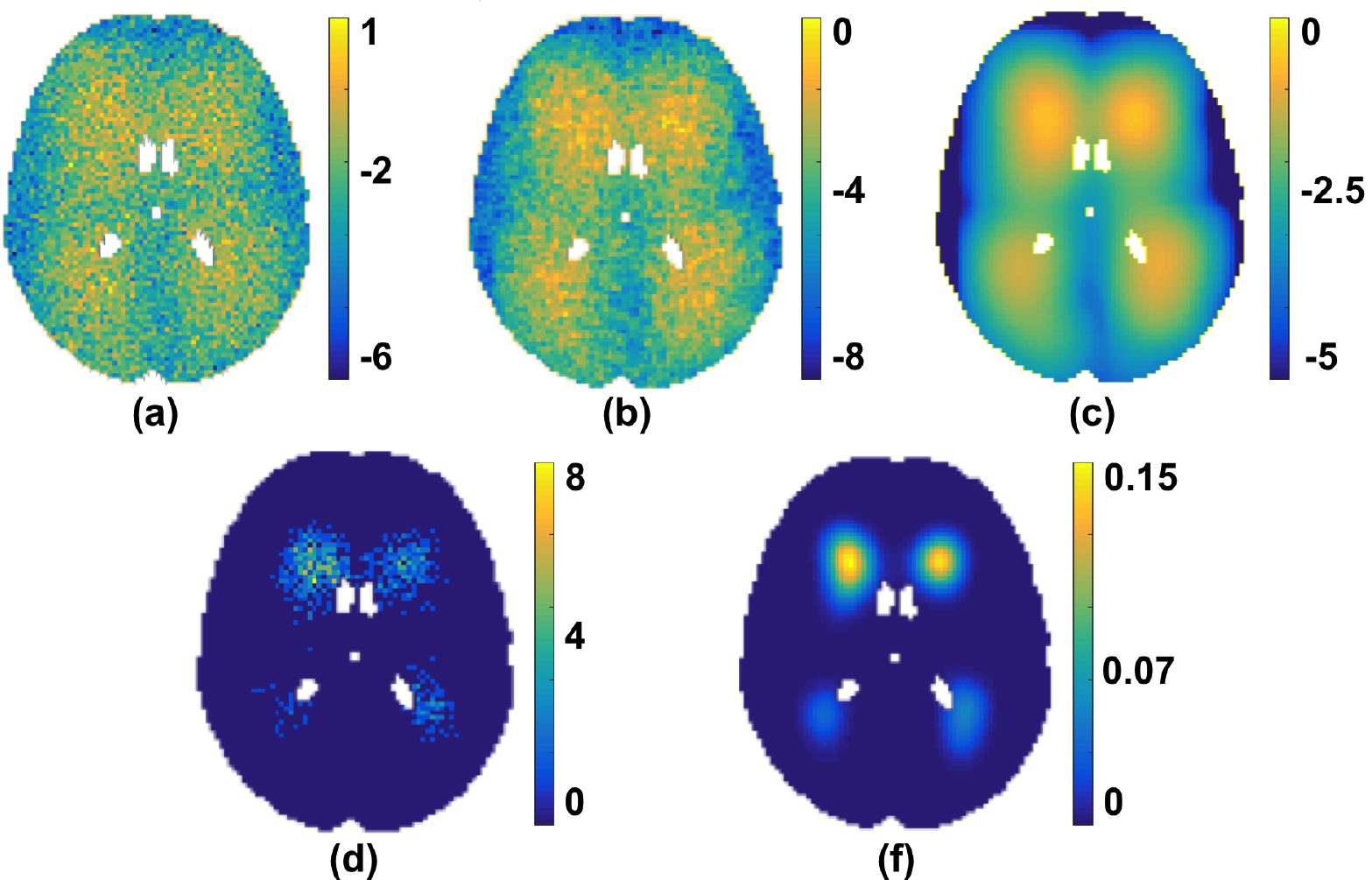
An instance of simulated lesion data (shown at slice *z* = 45 and age bin number *t* = 20). (a) Addition of random noise to *μ* indicating randomly sampled voxels, (b) lesion voxels after smoothing (a) with *K*, (c) *K* ∗ *μ*, (d) Sum of binary lesion maps (*R*_*sim*_) in a specific age group, (e) Simulated *θ*_*true*_ image

#### 2.2.1. Comparison with the existing alternative methods

We applied our algorithm to the simulated data and compared its performance with respect to some alternative methods with different levels of complexity. The simplest method for estimating the lesion probability with respect to age groups is to determine the group-level average of lesion probability maps as done in Rostrup et al. (2012). We performed the similar average on binary lesion maps and calculated *Average* = *R*_*t*_/*N*_*t*_ for each age group *age*_*t*_. However, the resulting lesion probability map *Average* is limited by the number of subjects in each *age*_*t*_ and does not provide a continuous/generalized estimate of lesion distribution. Hence another option is to smooth the average lesion distribution map *Average*, using a Gaussian kernel with different standard deviations *σ* (0.5, 1.5 and 3.0) to get a more spatio-temporally continuous lesion probability map *Smoothed_Average*(*σ*). We also applied the Bayesian Spatial Generalized Linear Mixed Model (BSGLMM) based on spatially varying coefficients proposed by Ge et al. (2014). We evaluated the performance of the above methods by determining the error values (difference between the ground truth probabilities and the outputs of the above methods) and the corresponding mean-squared error (MSE) values.

### 2.3. Application to real data

We applied our algorithm to two different groups from the UK Biobank data and compared the lesion distribution between the two groups using the methods that provided better MSE values than those reported in the literature, as determined from tests on the simulated dataset. We further validated our observations by applying our method on a clinical dataset consisting of a vascular population of subjects who had experienced a transient ischemic attack or minor stroke.

The dataset used for initial validation is a subset of UK Biobank data (Miller et al. (2016)). In the UK biobank data, about 10% of the sample has been identified as being diagnosed with hypertension, corresponding to around 7,975 subjects. Our goal was to model the lesion distribution with respect to age, comparing the subjects diagnosed with hypertension (HT group) to subjects without hypertension, where the latter acts like a control group (non-HT group). In this case, to reduce the amount of computation, we selected a balanced subset of randomly sampled 1000 subjects, 500 subjects from the HT group (age range 45.5 - 78.3, mean age 66.3 ± 6.1 years, with female to male ratio, F:M = 202:298), while the remaining were from the non-HT group (age range 45.5 - 78.4, mean age 62.0 ± 7.6 years, with female to male ratio, F:M = 257:243).

For these subjects we generated binary lesion masks to be used as input for our algorithm. This was performed using BIANCA (Griffanti et al. (2016)) on FLAIR images, also using information from T1-weighted images (Alfaro-Almagro et al. (2018)). The lesion probability maps obtained were binarised after thresholding at 0.8. We then registered the single subject binary lesion maps to the MNI space using linear and non-linear registration, thresholded them at 0.5 and binarised them to compensate for interpolation. We then applied our algorithm to the two groups separately and compared the results with those from other methods. When applying our method, we arranged the subjects in each group into one year age bins to form a 4D image of dimension 91 × 109 × 91 × 33. We compared the two groups using an unpaired t-test using permutation testing randomise, (Winkler et al. (2014)) to analyse the significance of the difference in the lesion probability values between HT and non-HT groups, and calculated the corresponding z-scores. Results were considered significant for *p*_*corr*_ < 0.05, corrected for multiple comparisons by using the null distribution of the max (across the image) voxelwise test statistic. For BSGLMM, the t-values were obtained by dividing the posterior mean by posterior standard deviation (Ge et al. (2014)).

Finally, we validated our algorithm on a clinical dataset of subjects at risk of vascular cognitive impairment. The dataset consists of MRI data from 474 consecutive eligible participants from Oxford Vascular Study (OXVASC) (Rothwell et al. (2004)), who had recently experienced a minor non-disabling stroke or transient ischemic attack (TIA) (age range 20 - 102 years, mean age 67.4 ± 14.3 years, with female to male ratio, F:M = 240:234) (see Griffanti et al. (2016) for more details). Similar to the UK Biobank dataset, we obtained the lesion binary masks using BIANCA and registered the single-subject masks to the MNI space. We grouped the data in age bins of 3 years to form a 4D image of dimensions 91 × 109 × 91 × 27 and applied our model. We also performed permutation-based statistical analysis (corrected for multiple comparisons) to test the statistical significance of the effect of age on the lesion probabilities. For this analysis, in each iteration, we permuted the data with respect to the age groups and applied our model on the permuted data to generate results for that permutation, that were then used to assess statistical significance, with correction for multiple comparisons (using the maximum statistic across space).

## 3. Results

### 3.1. Results on the simulated data

Figure 5 shows the error values of the outputs of the different methods with respect to ground truth, while Table 1 reports the corresponding MSE values and running time. For the *Smoothed_Average*(*σ*) method, we tried various values of standard deviations *σ* for the Gaussian kernel and have shown the results for the three *σ* values that provided minimum MSE values for this simulated dataset.

**Figure 5:**
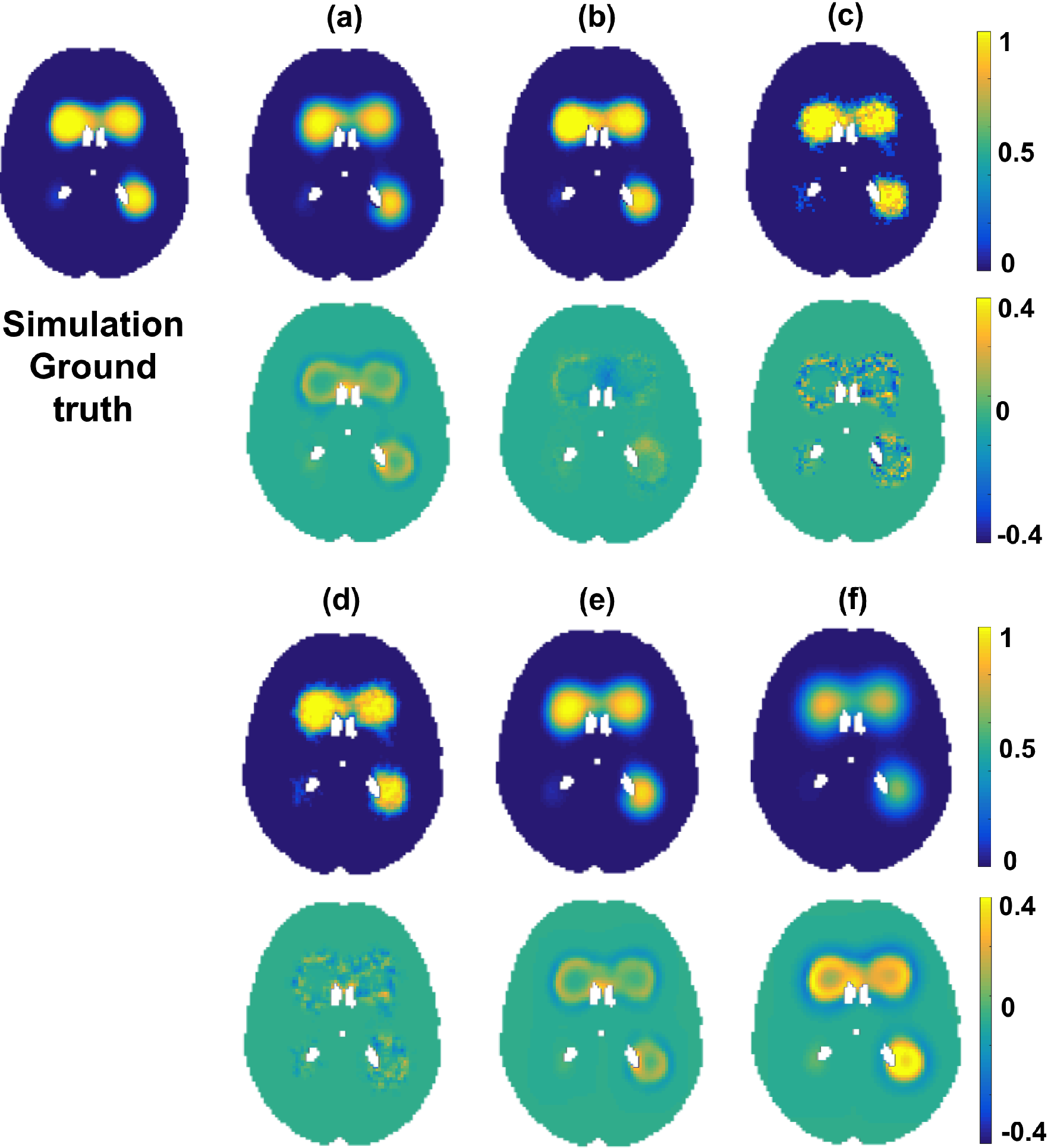
Comparison of simulation analysis results using different methods. The results are shown at slice *z* = 45 and age bin number *t* = 33. Ground truth distribution *θ*_*true*_ shown along with the outputs of various methods 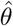 (a - f) and the corresponding error maps immediately underneath the probability maps. The outputs are shown for our Bayesian spline method (a), BSGLMM (b), *Average* (c), *Smoothed_Average*(*σ* = 0.5) (d), *Smoothed _Average*(*σ* = 1.5) (e) and *Smoothed_Average*(*σ* = 3) (f).

On a simulated dataset consisting of 698 subjects, we obtained an MSE value of 7.27 × 10^−5^ for our Bayesian spline method, while BSGLMM method gave a lower MSE value of 5.22 × 10^−5^ with ≈ 15 times slower running time. *Average* is the fastest (with < 1 second running time) but gives twice as much as error (18.790 × 10^−5^) as our method. Ge et al. (2014) reported an MSE value of 12 × 10^−5^ on simulated data consiting of 100 randomly sampled subjects. From Table 1, we can observe that only *Average* and *Smoothed_Average*(*σ* = 3) give MSE values greater than 12 × 10^−5^ (reported in Ge et al. (2014)). Moreover, both MSE values and running times increases as the standard deviation *σ* of the smoothing kernel increases due to increase in the kernel size.

**Table 1:** Comparison of MSE values and running time of different methods on simulation data

| Method | MSE values | Running time / Processor specs |
| --- | --- | --- |
| Our Bayesian spline method | $7.27 \times 10^{-5}$ | < 20 mins (84 secs per iteration) / iMac CPU with 2.9 GHz Intel Core i5 processor |
| BSGLMM (Ge et al. (2014)) | $5.22 \times 10^{-5}$ | $\approx 293$ mins / NVIDIA Tesla K80 GPU with 12GB |
| *Average* | $18.79 \times 10^{-5}$ | < 1 sec / iMac CPU with 2.9 GHz Intel Core i5 processor |
| *Smoothed_Average*($\sigma = 0.5$) | $3.86 \times 10^{-5}$ | $\approx 8$ secs / iMac CPU with 2.9 GHz Intel Core i5 processor |
| *Smoothed_Average*($\sigma = 1.5$) | $8.56 \times 10^{-5}$ | $\approx 11$ secs / iMac CPU with 2.9 GHz Intel Core i5 processor |
| *Smoothed_Average*($\sigma = 3$) | $33.71 \times 10^{-5}$ | $\approx 17$ secs / iMac CPU with 2.9 GHz Intel Core i5 processor |

Figure 6 shows the lesion probability with respect to age at a representative periventricular (a) and deep (b) voxel for the tested methods. It is worth noting that BSGLMM posterior probabilities tend to increase monotonically with age throughout the white matter irrespective of the location of the lesions.

**Figure 6:**
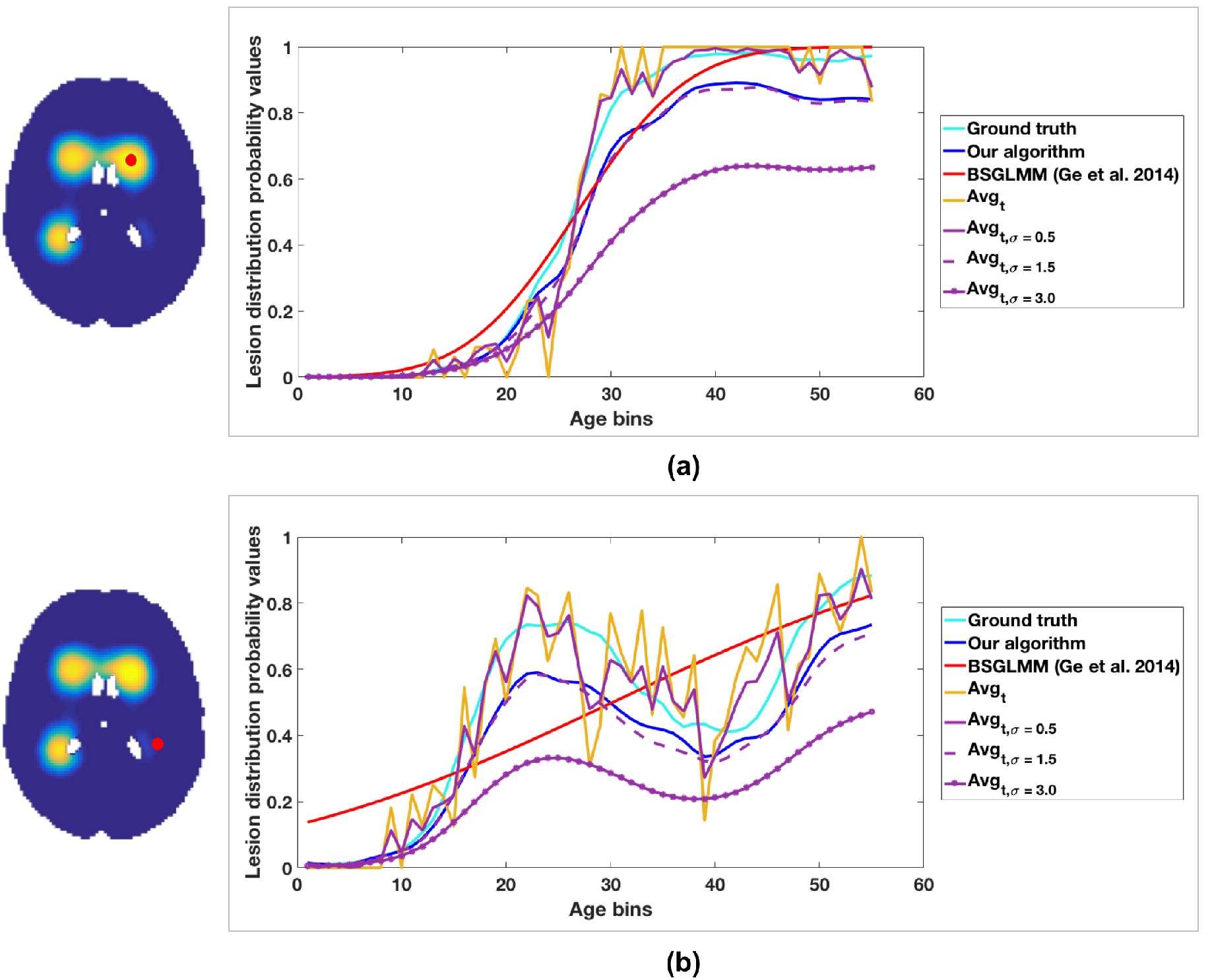
Lesion probability values plotted versus age. The plot of lesion probability values shown for a periventricular voxel (a) and a deep voxel (b). The red dot on the ground truth images indicate the voxel locations corresponding to the plot.

We also tested the effect of change in cubic b-spline knot spacing and the standard deviation of kernel *K* used in the ground truth estimation on the error values (see supplementary material). We observed that the cubic b-splines with knot spacing of 2 voxels provided the best result on the simulated data and hence we used the same specifications henceforth on the real data experiments.

### 3.2. Results on the real data

We show the results of our algorithm on UK Biobank data to estimate the lesion distribution probabilities with respect to age within the HT and non-HT subjects in Figure 7.

**Figure 7:**
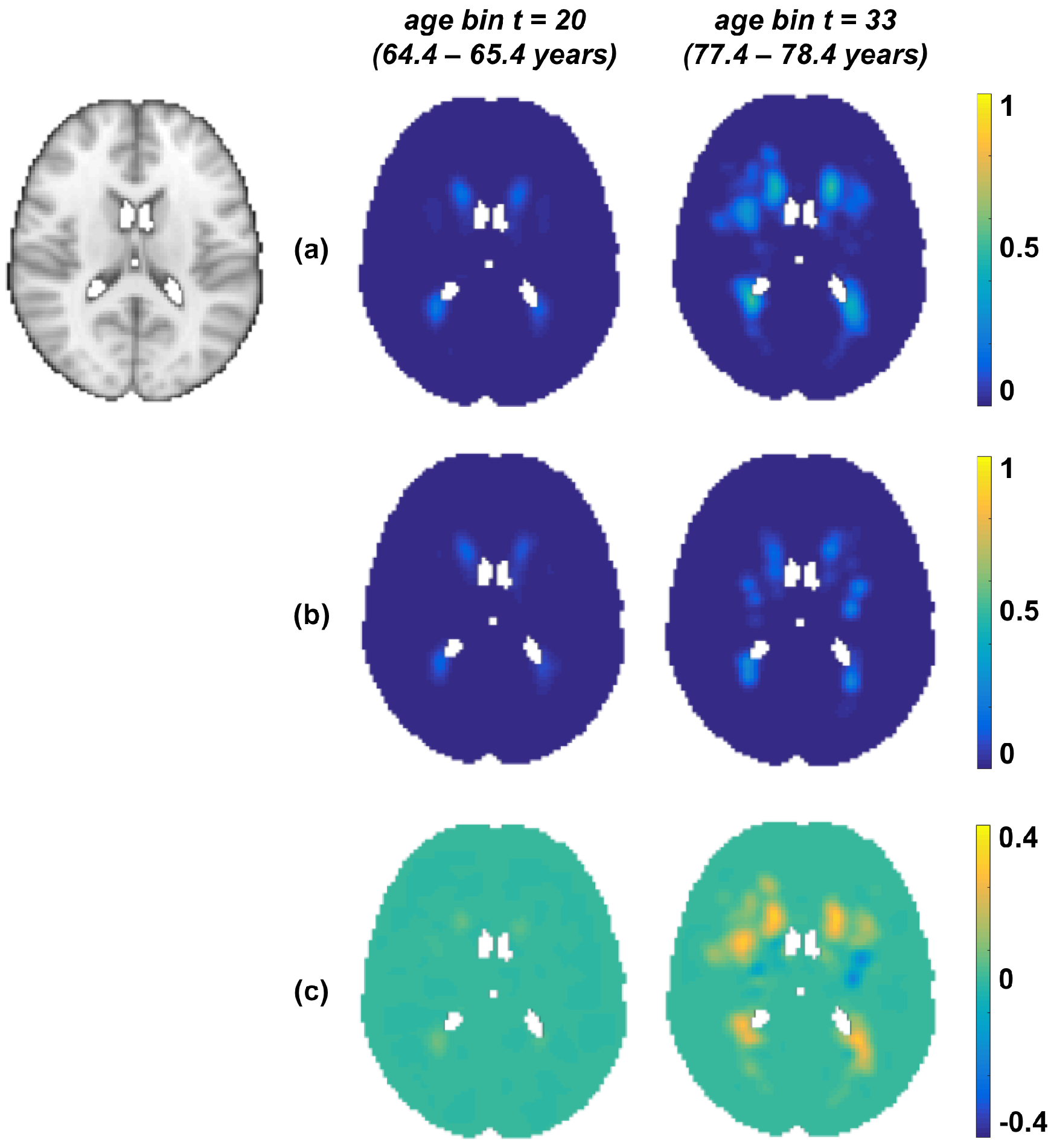
Results on UK Biobank data. Output of our Bayesian spline method on HT group (a) and non-HT group (b) shown along with their difference maps (c). First column shows the age group t = 20 (*age*_*t*_ = 64.4 - 65.4 years) and second column shows t = 33 (*age*_*t*_ = 77.4 - 78.4 years). All the results have been shown at the slice z = 45 (MNI template on the left)

The lesion probabilities increased with age for both the groups and are higher in the HT group (≈ 0.4 as shown in fig. 7(c)) than the non-HT group. We can observe that this difference is higher in the periventricular regions compared to deep regions, and higher in the older age groups (≈ 0.3 between the columns in row (c)) compared with the younger group. So not only is there a strong relationship between lesion probabilities and hypertension, but also this relationship is more pronounced as the population gets older.

In figure 8 and figure 9, we show the results of our algorithm compared with the posterior probabilities obtained from the BSGLMM method and the outputs of smoothed average maps (*Smoothed_Average*(*σ* = 0.5), *Smoothed_Average*(*σ* = 1.5)) respectively. We chose to compare the above methods with our Bayesian spline algorithm since these methods gave MSE values lesser than 12 × 10^−5^ (Ge et al. (2014)) on the simulated data (Table 1). We observed that the lesion probabilities in periventricular regions increased with age for both HT and non-HT groups, while the probabilities in deeper regions increased only in the HT group (between the first two rows in (a) in fig. 8 and 9). The bottom rows of figures 8 and 9 show the z-scores for significant regions (*p*_*corr*_ < 0.05), obtained from a statistical test to determine the significance of differences between the HT and non-HT groups. We observed increased lesion probabilities in the HT group compared with non-HT group, especially in the deep regions. This effect is particularly strong in our method and *Smoothed_Average*(*σ* = 1.5) when compared with BSGLMM and *Smoothed_Average*(*σ* = 0.5) (fig. 8 and 9 (bottom row)).

**Figure 8:**
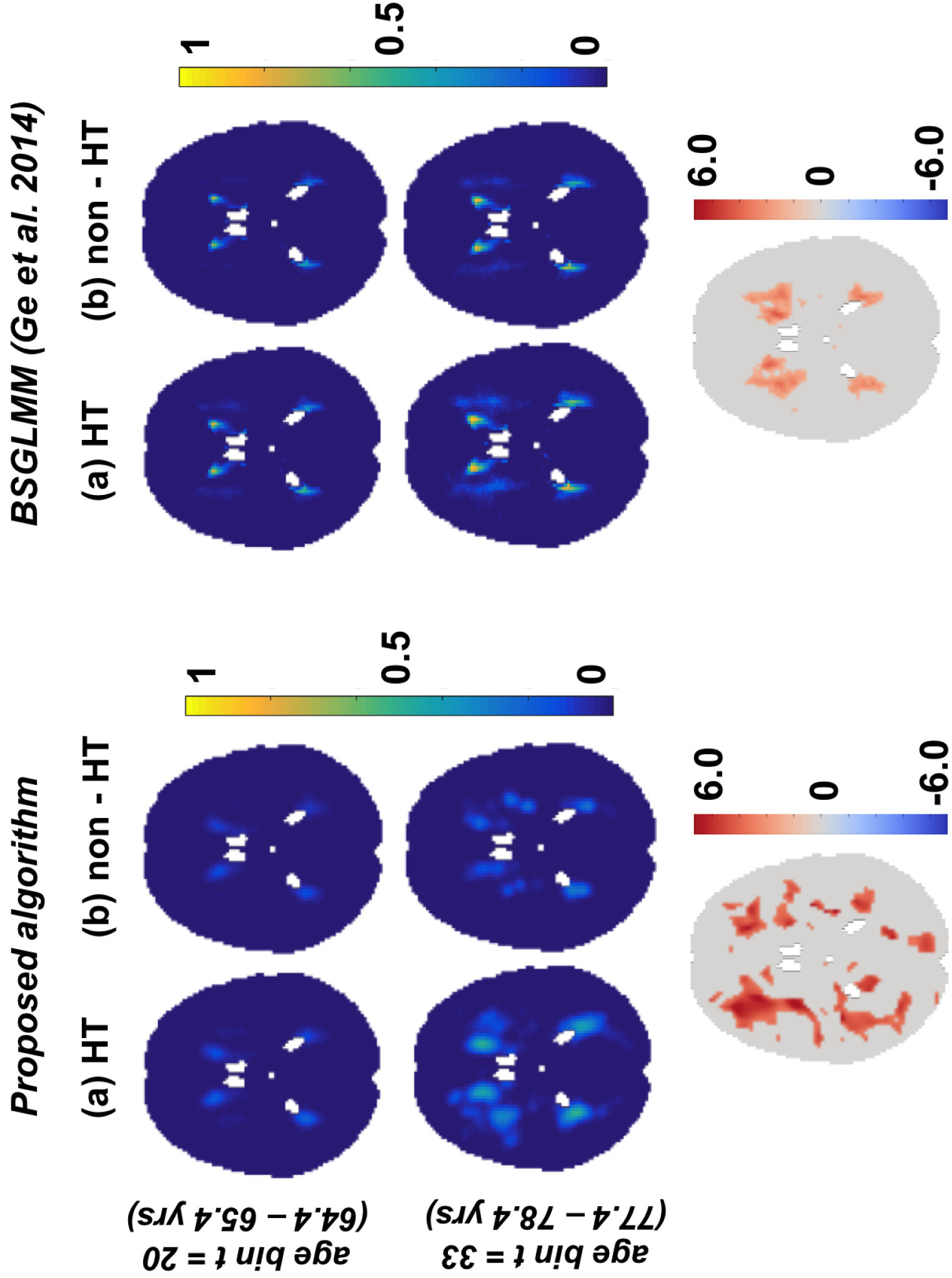
Comparison of the results of the Bayesian spline algorithm with BSGLMM on UK Biobank data. Top two rows show the results of the methods for HT and non-HT groups for two age groups. The bottom row shows the z-scores in the significant regions (*p*_*corr*_ < 0.05), obtained from the significance test on the differences of the lesion probabilities between HT and non-HT groups obtained by the corresponding methods.

**Figure 9:**
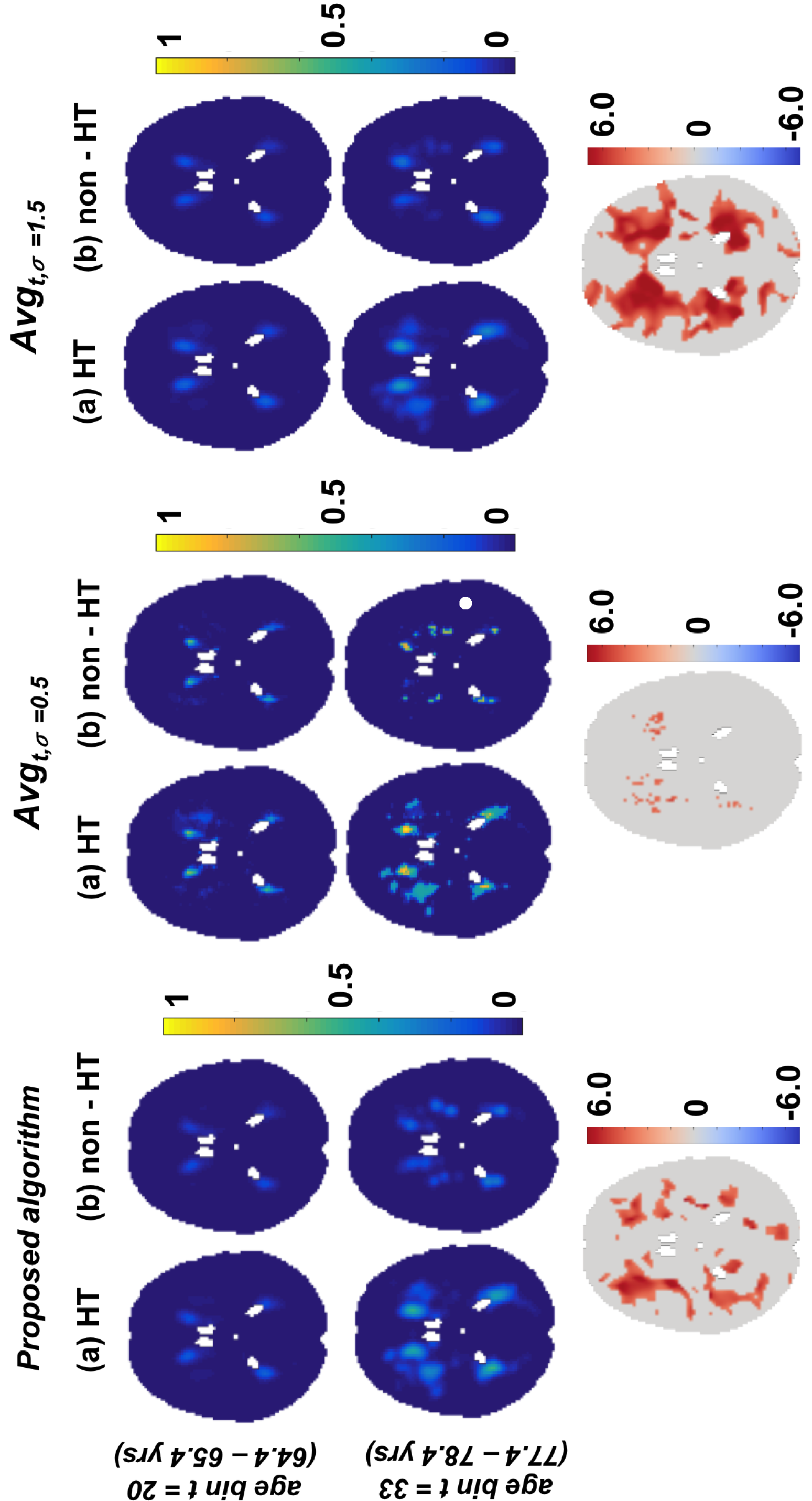
Comparison of the results of the Bayesian spline algorithm with *Smoothed_Average*(*σ* = 0.5) and *Smoothed_Average*(*σ* = 1.5) on UK Biobank data. Top two rows show the results of the methods for HT and non-HT groups for two age groups. The bottom row shows the z-scores in the significant regions (*p*_*corr*_ < 0.05), obtained from the significance test on the differences of the lesion probabilities between HT and non-HT groups obtained by the corresponding methods.

Figure 10 (top row) shows the estimated lesion probabilities from our Bayesian spline method on OXVASC data for two representative age groups. Similar to our results on UK Biobank data, it can be observed that the overall lesion probability values are relatively higher in the older age group compared to the younger age group (≈ 0.2, between second and third columns). On performing the permutation test, we observed that periventricular voxels change significantly with age (*p* < 0.05) as shown in Figure 10 (bottom row).

**Figure 10:**
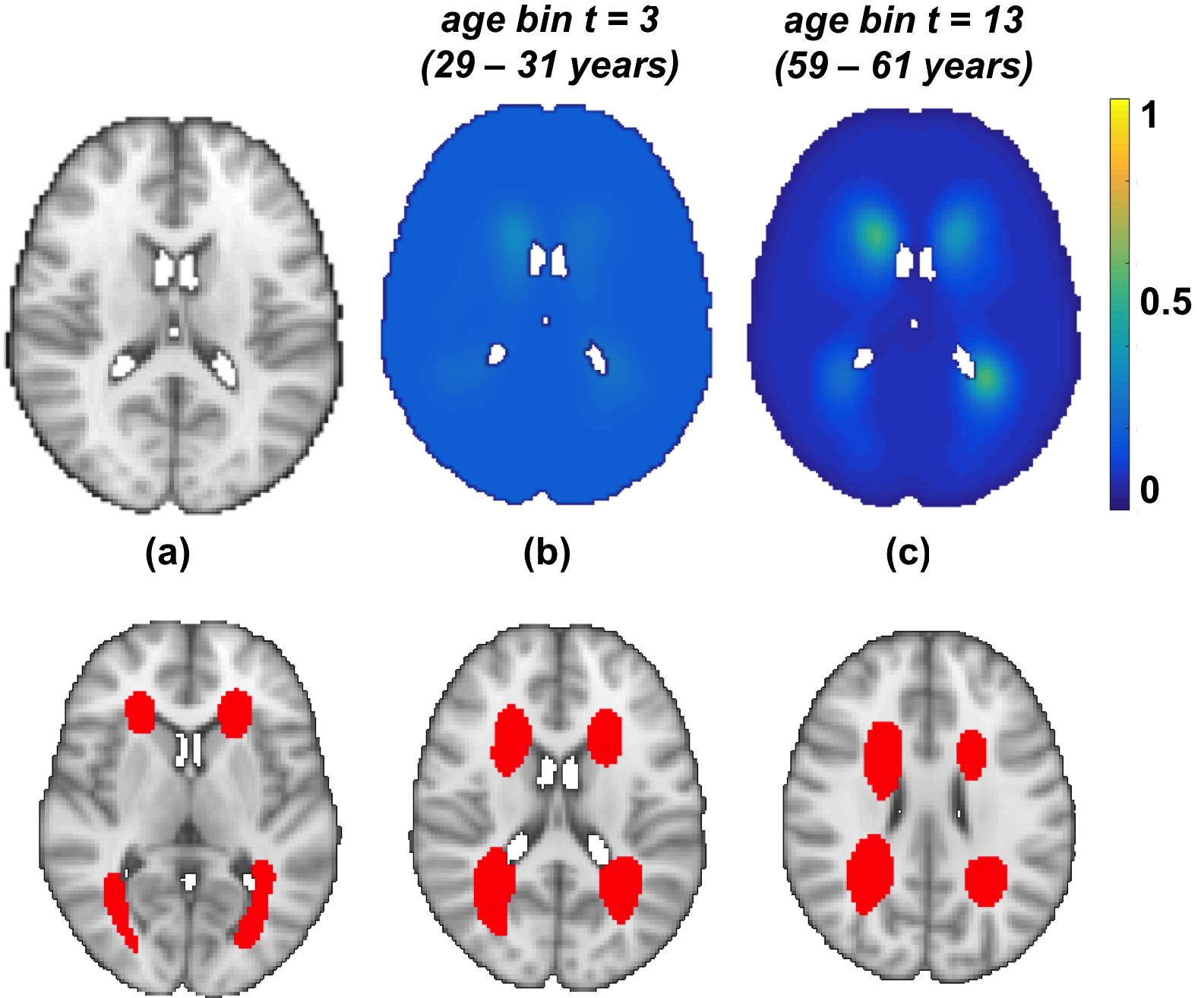
Results on OXVASC data. (Top row) From left to right: MNI brain, lesion probability distribution at bin numbers *t* = 3 (*age*_*t*_ = 29 - 31 years) and *t* =13 (*age*_*t*_ = 59 - 61 years). The results are shown at slice *z* = 45. (Bottom row) Results of permutation analysis. From left to right: Voxels that change significantly (*p* < 0.05) with age (red) shown on slices *z* = 39,45 and 49

## 4. Discussion and conclusion

In this work we compared several methods including our proposed Bayesian spline method to model the distribution of white matter lesions with respect to a parametric factor within different populations. For the experiments in this paper, we considered age as our parameter of interest, but the framework is general and can work with any continuous parameter of interest.

During the development, we analysed the effect of various parameters of our algorithm on the resulting lesion probabilities. One of the most important parameters in the algorithm is the knot spacing in the spline model. We studied its effect and how it relates to the size of smoothing kernel *K* used for the generation of the simulation data by varying them independently and evaluated their combined effect on the algorithm result 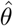 (see supplementary material). The best results were obtained for the knot spacing of 2 with MSE of 7.270 × 10^−5^. We also evaluated the effect of the initialization provided for the optimization function and observed that the algorithm converges quicker when we provide the smoothed average lesion map as the initial value for λ_*i*_, whereas it fails to converge when a zero image or unity image is provided as initialization for λ_*i*_. Also, our algorithm converges to the same output when the average lesion maps are smoothed with kernels of similar standard deviation.

When comparing the results of different methods on the simulated dataset, we found that the MSE values are lower than that reported in Ge et al. (2014) for most of the methods, except *Average* and *Smoothed_Average*(*σ* = 3.0). We obtained the highest MSE value and highest speed for the simplest method *Average*. However, this also gave a resulting image that was not as spatially continuous as the other methods. Moreover, the accuracy of *Average* (as well as of its Gaussian smoothed results *Smoothed_Average*(*σ*)) depends on the number of subjects in the age groups. While they provide reliable estimates in larger populations, their estimates are not so accurate when there are fewer subjects or when the lesion load is lower, since the data becomes sparse. Also, in the case of smoothed *Smoothed_Average*(*σ*) method, determining the optimum value of standard deviations *σ* for the Gaussian kernel is dependent on the dataset and the lesion characteristics. However, the smoothed *Smoothed_Average*(*σ*) outputs provide good initial estimates for our method and lead to better convergence, as we discussed above.

We observed that, while results of both the Bayesian spline method and BSGLMM show relatively similar MSE values, the spline modelling in our algorithm is spatially more robust when compared to BSGLMM method. For instance, from Figure 6, while lesion probabilities obtained from BSGLMM method fit well in periventricular areas, they do not fit the ground truth distribution as well in deep areas (figure 6). Another important advantage of our method is that it is much faster than BSGLMM (Table 1) and can easily scale up to very large numbers of subjects. Our modelling algorithm takes less than 20 minutes (84 seconds per iteration), when run on an iMac with 2.9 GHz Intel Core i5 processor. This is due to the fact that we independently perform the convolution in all 4 dimensions with a 1D spline which reduces the computational load. Also, by calculating the analytical gradient (Eqn. 16) we avoid the numerical computation of gradient for 4D data in the optimization step. Even though the maximum number of iterations allowed for the optimization step is 50, our algorithm typically converges earlier (10 iterations on simulated data and < 10 on real data). On the other hand, the BSGLMM algorithm is computationally demanding since it has to estimate the posterior distribution via Markov Chain Monte Carlo model using Gibbs sampler at each voxel.

Our results on the real data (UK Biobank and OXVASC datasets) show that the lesion distribution probabilities are higher in the later ages, which is in line with the existing literature (Simoni et al. (2012), Alfaro-Almagro et al. (2018)). Moreover, the fact that we observed an increase in lesion probabilities in deep white matter regions in the HT group with respect to the non-HT group for all the methods (fig. 8 and 9), suggests that deep lesions might be associated with hypertension. The same phenomenon has also been reported in Rostrup et al. (2012). We further verified it with our statistical tests and observed higher value of z-scores with greater significance (*p* < 0.05), particularly in the deep regions between the HT and non-HT groups. Though this is generally observable in all the methods, it is more evident for for our Bayesian spline method and smoothed average map *Smoothed_Average*(*σ* = 1.5) i.e. deep regions are affected significantly in the HT group compared with the non-HT group for these two methods.

The results on the UK Biobank data are similar to those on the simulated data when comparing the Bayesian spline method with BSGLMM and *Smoothed_Average*(*σ* = 0.5). In particular, we found that the results of our method appear smoother by comparison to the other two. In the case of *Smoothed_Average*(*σ* = 0.5), although the method gave MSE value lower than our method on simulated data, on a real dataset like UK biobank it did not yield a smooth lesion distribution and had isolated voxels in the lesion probability maps. This is because the method is affected by the sparsity of the real data and hence is dependent on data/lesion characteristics. Also, statistical test result of *Smoothed_Average*(*σ* = 0.5) shows only a few isolated voxels that are affected significantly more in the HT group, when compared with the non-HT group.

To conclude, we compared different methods to model the distribution of white matter hyperintensities within a population with respect to a parametric factor of interest, such as age. The evaluation of the proposed Bayesian spline method on the simulated data provides an MSE value of 7.27 × 10^−5^, with the advantage of being computationally more efficient. We compared the results of various methods on a real dataset between hypertension and control groups and found that deep lesions increase significantly with hypertension, which is inline with existing work. We validated our model on real data showing that periventricular lesions increase significantly with age, which is well established in the literature. The results are consistent between the two datasets, showing that our model is robust across datasets. As future work, the Bayesian spline method could be used to model the distribution of lesions with respect to other factors. In addition, the lesion probability distribution map obtained from our model can be used either as an additional feature or as a spatial prior to improve lesion segmentation algorithms.

## Acknowledgements

We acknowledge all the participants. We acknowledge the use of the facilities of the Acute Vascular Imaging Centre, Oxford. The data used in this work was obtained from UK Biobank under Data Access Application 8107. We are grateful to UK Biobank for making the resource data available, and are extremely grateful to all UK Biobank study participants, who generously donated their time to make this resource possible.

The authors report no biomedical financial interests or potential conflicts of interest.

